# A2 and A1B *in vitro* milk digests: effects on *in vitro* leaky gut model and adipose cells

**DOI:** 10.64898/2026.05.09.723973

**Authors:** Jessica Perugini, Paola Bendinelli, Edoardo Scopini, Chiara Galli, Stefano Cattaneo, Valentina Bonfatti, Saverio Cinti, Adele Finco, Ivano De Noni, Antonio Giordano, Anita Ferraretto

## Abstract

Obesity is associated with chronic low-grade systemic inflammation of adipose tissue and is often linked to intestinal epithelial barrier (IEB) dysfunction. The present study aimed to evaluate the effects of *in vitro* gastrointestinal digests of bovine milk containing A1B or A2 β-casein variants on leaky IEB and adipocyte inflammation. Digests of A1B (DA1B) and A2 (DA2) milk were administered to an *in vitro* Caco-2/HT-29 intestinal cell co-culture mimicking a leaky gut. Intestinal absorbed fractions derived from A1B (MA1B) and A2 (MA2) were administered to hMADS adipocytes. DA1B and DA2 did not modify intestinal permeability, either in the absence or the presence of inflammation. DA1B reduced Claudin-1 mRNA, as well as zonula occludens-1 mRNA and protein expression. Both DA1B and DA2 increased interleukin-8 expression, but only DA1B increased tumor necrosis factor-α. In human adipocytes, MA1B, and to a lesser extent MA2, increased the expression of pro-inflammatory markers monocyte chemoattractant protein-1 and interleukin-6, while reducing adiponectin levels. DA2 preserved *in vitro* leaky IEB integrity and exhibited a lower inflammatory potential in both leaky gut and adipocytes compared to DA1B. This study is the first to establish a link among A2 milk, leaky gut syndrome, and obesity.

## 1. Introduction

Obesity represents a chronic low-grade inflammatory state driven by excessive secretion of proinflammatory cytokines from adipose tissue, which has been increasingly associated with intestinal barrier dysfunction [1, 2]. In individuals with obesity, chronic lipid and carbohydrate overload results in adipocyte hypertrophy, stress, and death, all conditions that induce tissue inflammation characterized by adipokine dysregulation, fibrosis, and infiltration by inflammatory cells, mainly macrophages [3, 4]. These phenomena are accompanied by abnormal release of fatty acids (FA), adipokines, and proinflammatory molecules, including tumor necrosis factor-alpha (TNFα), interleukin-6 (IL6), and monocyte chemoattractant protein-1 (MCP1), which induce insulin resistance and favor the onset of type 2 diabetes by acting on target organs [5]. At the same time, diet composition, both in terms of macronutrient intake and specific dietary components, is known to display several therapeutic effects, for example, crucial anti-inflammatory properties for the prevention of obesity [6]. Among foods, bovine milk proteins, when submitted to gastrointestinal digestion, represent an important source of essential amino acids and peptides displaying biological functions and regulatory properties [7, 8]. In bovine milk, β-casein (β-CN) may exist in different genetic variants, with the A1, A2, and B being the most common isoforms. These variants differ in the amino acid residue occurring at position 67 of their primary sequences. A histidine residue is present in A1 and B variants, while it is replaced by proline in the A2 variant [9]. This substitution decreases the likelihood of releasing the bioactive peptide β-casomorphin-7 (BCM-7) during gastrointestinal digestion of A1 and B variants. This peptide can bind to µ-opioid receptors, affecting gastrointestinal physiology as well as the cardiovascular, neurological, and endocrine systems [10]. Based on these observations, the consumption of A1 milk has been negatively associated with human health, whereas A2 milk was hypothesized to be protective [11]. The B variant shares the same biochemical feature as A1, but its relationship with human health has not been directly investigated. Although preclinical and clinical studies have failed to demonstrate a real benefit from A2 milk toward cardiovascular and neurological disorders, some evidence suggests potential beneficial effects on gut health [12-14], especially in presence of a leaky gut characterized by a low-grade inflammation and dysbiosis, a situation mainly observed in celiac and lactose intolerant individuals, intestinal bowel diseases [15] as well as gut aging [16, 17], and in the presence of obesity [18].

Studies conducted on humans and animals affected by obesity have shown that the consumption of high levels of fats and carbohydrates leads to the leakiness of tight junction proteins in the intestinal cells and, as a consequence, to higher permeability to lipopolysaccharides and endotoxins. These, in turn, can reach the portal veins, causing endotoxemia associated with changed microbiota composition and bacterial translocation [19]. Based on this evidence, the maintenance and/or restoration of the intestinal barrier integrity could be one of the possible factors able to positively affect the onset of obesity, its progression, and/or to mitigate the associated metabolic consequences. Since obesity is a chronic inflammatory condition affecting adipose tissue, the intake of specific milk-derived macronutrients may play a crucial role in modulating its pathophysiology. However, the effects of A2 milk on adipocytes have never been investigated. Therefore, this study aimed to evaluate the effects of *in vitro* gastrointestinal digests from bovine A2 milk and from milk containing both A1 and B variants (A1B milk) on intestinal barrier integrity and inflammatory markers in an *in vitro* model of leaky gut, as well as the effects of their intestinal digested and absorbed fractions on inflammatory responses in adipocytes.

## 2. Materials and Methods

### 2.1. Materials

Reagents to cultivate intestinal cells were purchased from Merck (Milano, Italy) except for Fetal Bovine Serum (FBS), which was from EuroClone Ltd, West Yorkshire, UK. Reagents to cultivate adipocytes were purchased from Pan-Biotech GmbH (Aidenbach, Germany). Recombinant human Tumor Necrosis Factorα (TNFα), Interferonγ (IFNγ), and fibroblast growth factor (hFGF)-2 were from Peprotech (Cranbury, NJ, U.S.A.). Any other reagent, unless indicated, was from Merck.

### 2.2. Milk collection and analysis

Milk samples were kindly provided by the farm Angolo di Paradiso (Fermo, Italy) as part of the project I-Milka 2 - PSR MARCHE 2014. Details on the animals used, the creation of the experimental milk pool, sample collection, and analysis were described elsewhere [20]. Briefly, during a preliminary herd analysis, the β-CN milk phenotype of all the cows present on the farm was determined through RP-HPLC analysis according to the method developed and validated by Bonfatti et al. [21]. For each cow, the information on β-CN phenotype was coupled with milk composition data from the monthly herd recording. Between June and July 2020, on three occasions at two-week intervals, small groups of cows were selected and separately milked to create distinct milk pools for comparative analysis. On each of the three days, two milk pools were formed: one including β-CN A1 and B variants (A1B milk) and one including only the β-CN A2 variant (A2 milk).

To maximize the difference in the proportions of β-CN genetic variants between the two pools, while maintaining similar protein and fat levels, κ-casein (κ-CN) and β-lactoglobulin (β-LG) genetic variants, and comparable somatic cell count (SCC), a small number of cows (n=2–3) per pool were selected each day. On each of the three days, the chosen cows were separately milked, and their daily milk output was pooled into two refrigerated tanks and stored at 4°C for 24 h until milk sample collection. Each cow was sampled once. On the day of the milk pooling, milk yield records and SCC values were automatically collected for each cow using the DeLaval VMSTM V300 milking system (DeLaval, San Donato Milanese, Italy) equipped with DelProTM FarmManager software (version 5.11, DeLaval, San Donato Milanese, Italy). The cows selected for the milk pools produced between 24 and 36 kg/d of milk, differed in parity (from 1 to 4), days in lactation (from 161 to 304), and had SCC below 128,000 cells/mL. On each milk pool, after 24 h from milking, pH was measured with a pH-meter (HI98191, Hanna Instruments, Villafranca Padovana, Italy), whereas titratable acidity was measured with 0.25 mol/L NaOH according to the Soxhlet-Henkel method. Two 50 mL samples of milk were then collected from each tank, frozen at −20°C, and transferred to the laboratories of the University of Padova for the analysis of proximate composition, mineral profile, and protein profile. Milk analyses were described in detail elsewhere [20]. Briefly, dry-matter was determined by vacuum oven at 102°C, fat content was obtained by the Röse-Gottlieb Method, the total and casein nitrogen by the Kjeldahl method, and mineral content by the gravimetric method. Quantification of Ca, P, Mg, K, and Na in milk samples was performed through inductively coupled plasma optical emission spectrometry (ICP-OES) after acidic digestion. Analysis of protein composition was carried out by RP-HPLC, enabling the identification and quantification of genetic variants of caseins and whey proteins [21].

### 2.3. *In vitro* static gastrointestinal digestion (SGID) of A1B and A2 milk samples

The *in vitro* SGID of A1B and A2 milk samples was performed according to the INFOGEST protocol [22]. Simulated salivary (SSF), gastric (SGF), and duodenal (SDF) fluids were prepared accordingly. In detail, the milk sample (5 mL) was added with 5 mL of SSF at pH 7.0 and then supplemented with 10 mL of SGF-containing porcine pepsin (1000 U/mL of SGF) and rabbit gastric lipase (30 U/mL of SGF). The gastric digestion step was performed at 37°C for 2 h at pH 3.0 (adjusted with 1 mol/L HCl). Subsequently, 20 mL of SDF and bile salts (10 mmol/L) were added to the digest. The enzymes used for intestinal digestion were porcine trypsin (200 U/mL SDF), bovine chymotrypsin (50 U/mL SDF), porcine intestinal lipase (1000 U/mL SDF), and colipase (lipase/colipase at 1:2 molar ratio). The intestinal phase was conducted at 37°C for 2 h at pH 7.0 (1 mol/L NaOH). Each milk sample was subjected to three replicate digestions on the same day. Control digestions without milk were also performed. The A1B and A2 milk *in vitro* digests (DA1B and DA2, respectively) were immediately frozen, then lyophilized and stored at −40°C.

### 2.4. Leaky Caco-2/HT-29 70/30 cell co-culture setting and treatment with DA1B and DA2

Caco-2 (BS TCL 87) and HT-29 (BS TCL 132) colon carcinoma cell lines were purchased from Istituto Zooprofilattico Sperimentale (Brescia, Italy). Cells were grown separately in Roswell Park Memorial Institute 1640 (RPMI 1640) medium supplemented with 10% heat-inactivated Fetal Bovine Serum (FBS), 2 mol/L l-Glutamine, 0.1 mg/L streptomycin, 100,000 U/L penicillin, 0.25 mg/L amphotericin B, containing 13.9 mmol/L glucose in a 37°C humidified atmosphere with 5% CO2.

The *in vitro* model of leaky gut was obtained by a co-culture constituted by a mixture of differentiated 70% Caco-2 and 30% HT-29 cells sub-cultivated for more than 40 passages (Caco-2) and 20 passages (HT-29) in complete RPMI 1640 medium [23]. Under these experimental conditions, the co-culture spontaneously is endowed with transepithelial electrical resistance (TEER) values below 50 Ω cm^2^, unlike what is reported for a healthy human small intestine, which shows TEER values of 50-100 Ω cm^2^ [24]. The leaky co-culture was plated at a 40,000 cells /cm^2^ density and maintained for 10 d (6 d after confluence) before all the experiments. The quantity of DA1B and DA2 to be administered to the leaky co-culture as µg of digestate/cm^2^ of cell growing area was calculated based on the average daily intake of 250 g of milk, as recommended [25], and their distribution over an area of approximately 32 m^2^ of the human small intestine.

### 2.5. Cytotoxicity, TEER, and permeability of leaky co-culture

Co-culture was grown in 24 multiwell plates and treated with or without DA1B or DA2 diluted in the growth medium for 24 h. Cell viability, expressed as the ratio of live cells/total cells, was determined using an automatic cell counter, EVE™ NanoEntek (VWR International, Milano, Italy). Assays were performed at least in triplicate.

To measure TEER, the leaky co-culture was seeded on Transwell® Millicell® 24 insert plates (1.0 µm) assembled onto a Millicell® 24 well receiver tray (Merck) and treated for 24h with DA1B and DA2 diluted in the growth medium and administered on the apical side. DA1B and DA2 were tested before and after the IEB damage induced by Lipopolysaccharide (LPS, 100 µg/mL) administered in the apical side, and by a mixture (MIX) of TNFα (100 ng/mL) and IFN-γ (100 ng/mL) administered in the basolateral side. TEER was measured with a Millicell ERS system (Millipore Corporation, Burlington, MA, USA) at 37°C after 24 h incubation. At least three wells for each sample were set up, TEER was measured at three different points of each well, and the results were expressed as Ω cm^2^.

The paracellular permeability assay was performed after TEER measurement. A solution of 100 µmol/L Lucifer Yellow (LY, a fluorescent probe) in HBSS/5 mmol/L HEPES was added to the apical chamber of each well. Apical and basolateral solutions were separately collected after 2 h, and the fluorescence was measured (excitation wavelength 475 nm, emission wavelength 540 nm) by GloMax Discover (Promega Corporation, Madison, WI, USA). LY concentration was assessed using a standard curve.

### 2.6. Cell-derived digested and absorbed fractions from A1B and A2 milk digests

To mimic what happens *in vivo* after the digestion, i.e., the absorption of digests and their passage to the bloodstream for reaching target organs, DA1B and DA2 diluted in HBSS 5 mmol/L HEPES to a final volume of 400 µL were administered to the apical side of transwells, where leaky co-culture was seeded. After 2 h of incubation, the content of the basolateral chamber (800 µL), i.e. the digested and absorbed fractions derived from A1B and A2 milk digests (MA1B and MA2, respectively), were collected.

### 2.7. Administration of MA1B and MA2 to hMADS adipocytes

Human multipotent adipose-derived stem cells (hMADS), kindly provided by Dr Christian Dani (Université Côte D’Azur, Nice, France), were cultured as described previously [26, 27]. In brief, hMADS grown in low-glucose (1 g/L) proliferation medium (Dulbecco’s modified Eagle’s medium [DMEM]) supplemented with 10% FBS and 2.5 ng/mL hFGF-2 were used between the 16th and the 19th passage. To induce adipose differentiation, they were seeded in proliferation medium on multi-well plates at a density of 4,500 cells/cm^2^. When they reached the confluence, hFGF-2 was not replaced. The next day (day 0), cells were incubated in an adipogenic medium (serum-free proliferation medium/Ham’s F-12 medium) containing 10 µg/mL transferrin, 5 µg/mL insulin, 0.2 nmol/L triiodothyronine, 100 µmol/L 3-isobutyl-1-methylxanthine, 1 µmol/L dexamethasone, and 100 nmol/L rosiglitazone. Dexamethasone and 3-isobutyl-1-methylxanthine were not replaced from day 3, and rosiglitazone from day 9. Cell lipid content was assessed by Oil Red O staining. Treatments and biological assays were carried out on differentiated hMADS adipocytes from day 12 to day 15. hMADS adipocytes seeded in 6 multiwell plates were exposed to 800 µL of MA1B or MA2 for 24h and then used for RNA extraction and detection of cytokine release.

### 2.8. Quantitative Reverse Transcription Polymerase Chain Reaction (qRT-PCR)

Total RNA from leaky co-culture cells treated with DA1B and DA2 for 24h or hMADS adipocytes treated with MA1B and MA2 for 24h was extracted with TRIZOL reagent (Invitrogen, Carlsbad, CA), purified, digested with ribonuclease-free deoxyribonuclease, and concentrated using RNeasy Micro kit (Qiagen, Milano, Italy) according to the manufacturer’s instructions. For determination of mRNA levels, 1 µg RNA was reverse-transcribed with High-Capacity cDNA RT Kit with RNase Inhibitor (Applied BioSystems, Foster City, CA) in a total volume of 20 µL. The qRT-PCR was performed using TaqMan Gene Expression Assays and Master Mix TaqMan (Applied BioSystems). All probes were from Applied BioSystems (Table 1). Reactions were carried out in a Step One Plus Real Time PCR system (Applied BioSystems) using 50 ng cDNA in a final reaction volume of 10 µL. The thermal cycle protocol included initial incubation at 95°C for 10 min followed by 40 cycles of 95°C for 15 s and 60°C for 20 s. A control reaction without reverse transcriptase in the amplification mixture was performed for each sample to rule out genomic contamination. Samples not containing the template were included in all experiments as negative controls. TATA box-binding protein (TBP) was used as an endogenous control to normalize gene expression. Relative mRNA expression was determined by the ΔCt method (2^-ΔCt^).

**Table 1.** Taqman probes all from Applied Biosystems #4453320.

| Target Gene | Assay ID |
| --- | --- |
| ADIPOQ | Hs00605917_m1 |
| CLDN1 | Hs00221623_m1 |
| IL6 | Hs00174131_m1 |
| IL8 | Hs00174103_m1 |
| MCP1 | Hs00234140_m1 |
| OCLN | Hs01049881_m1 |
| TBP | Hs00427620_m1 |
| TJP1 | Hs01551867_m1 |
| TNFα | Hs00174128_m1 |

### 2.9. Cytokine release measurement

Cytokine release was measured by ELISA in conditioned media collected at the end of the incubation of leaky co-culture with DA1B and DA2 and of hMADS adipocytes with MA1B and MA2. The media were centrifuged and stored at −80°C until analysis. Interleukin-8 (IL8) and IL6 concentrations (pg/mL) were measured according to the manufacturer’s instructions using Quantikine ELISA Human IL8 kit (#HS800, R&D Systems, Minneapolis, USA) and Human IL6 ELISA kit (501030, Cayman Chemical, Ann Arbor, MI) in leaky co-culture and hMADS adipocytes, respectively.

### 2.10. Western blotting

Leaky co-culture was treated with DA1B and DA2 for 24 h. Cell lysates were obtained by using lysis buffer containing 50 mmol/L Tris–HCl (pH 7.4), 1% NP-40, 1 mol/L EDTA, 150 mmol/L NaCl, 1 mmol/L sodium orthovanadate, 0.5% sodium deoxycholate, 0.1% SDS, 2 mmol/L phenylmethylsulfonylfluoride, and 50 mg/mL aprotinin. Samples were centrifuged, and protein concentrations were determined by the Bradford Protein Assay (Bio-Rad Laboratories, Segrate, Italy). Proteins were separated by SDS-PAGE and then transferred to a nitrocellulose membrane using the Trans-Blot TurboTM Transfer system (Bio-Rad). To check loading and transfer efficiency, membranes were visualized with Ponceau S solution (Santa Cruz Biotechnology, Santa Cruz, CA, USA). Membranes were then blocked for 1 h at room temperature (RT) in TBS-Tween-20 (50 mmol/L Tris-HCL [pH 7.6], 200 mmol/L NaCl, and 0.1% Tween-20) containing 5% non-fat dried milk and subsequently incubated overnight at 4°C with the primary antibody rabbit anti-TJP1 (Abcam, catalogue no. ab59720, 1:50 dilution). After washing in TBS-Tween-20 and incubation with the secondary antibody for 1 h at RT (Jackson, catalogue no. 715-036-150, 1:5,000 dilution), bands were visualized with the Chemidoc Imaging system using the Clarity™ Western ECL chemiluminescent substrate (all from Bio-Rad). Quantitation of immunoreactive bands was performed using the Bio-Rad Image Lab software. Where appropriate, membranes were stripped, washed, and re-probed for total protein content.

### 2.11. Statistical analysis

Differences between A1B and A2 milk in pH, titratable acidity, mineral content (Ca, P, Mg, K, and Na), milk composition (total solids, fat, protein, casein, casein index, and lactose), and protein composition (such as casein ratios and specific casein components) were assessed using the Kruskal-Wallis test. This test was selected due to the small sample size (n = 3), which precluded the validation of normality assumptions for the distributions of the variables. The analysis was performed separately for each variable, and statistical significance was set at p < 0.05. Statistical analyses were conducted using R software (version 4.4.1).

All experiments with leaky co-culture and hMADS adipocytes were performed at least in triplicate. Comparisons were performed with one-way analysis of variance (ANOVA) followed by Tukey’s multiple comparisons post-hoc test. Results are reported as mean ± standard deviation (SD) or mean ± standard error of the mean (SEM), as indicated in the figure legends. A p-value < 0.05 was considered significant. All statistical analyses were performed with GraphPad Prism 10.4.2 software.

## 3. Results

### 3.1 Composition of A1B and A2 milk samples

The RP-HPLC enabled the identification and quantification of β-CN genetic variants as well as the other major casein and whey protein fractions. Chromatograms from A1B and A2 milk samples reported in Figure 1 clearly differentiated milk samples. Indeed, A1B milk displayed peaks corresponding to β-CN A1 and β-CN B variants, whereas only the A2 peak characterized the A2 milk. The chromatograms also allowed the identification and relative quantification of κ-CN and β-LG genetic variants.

**Figure 1.**
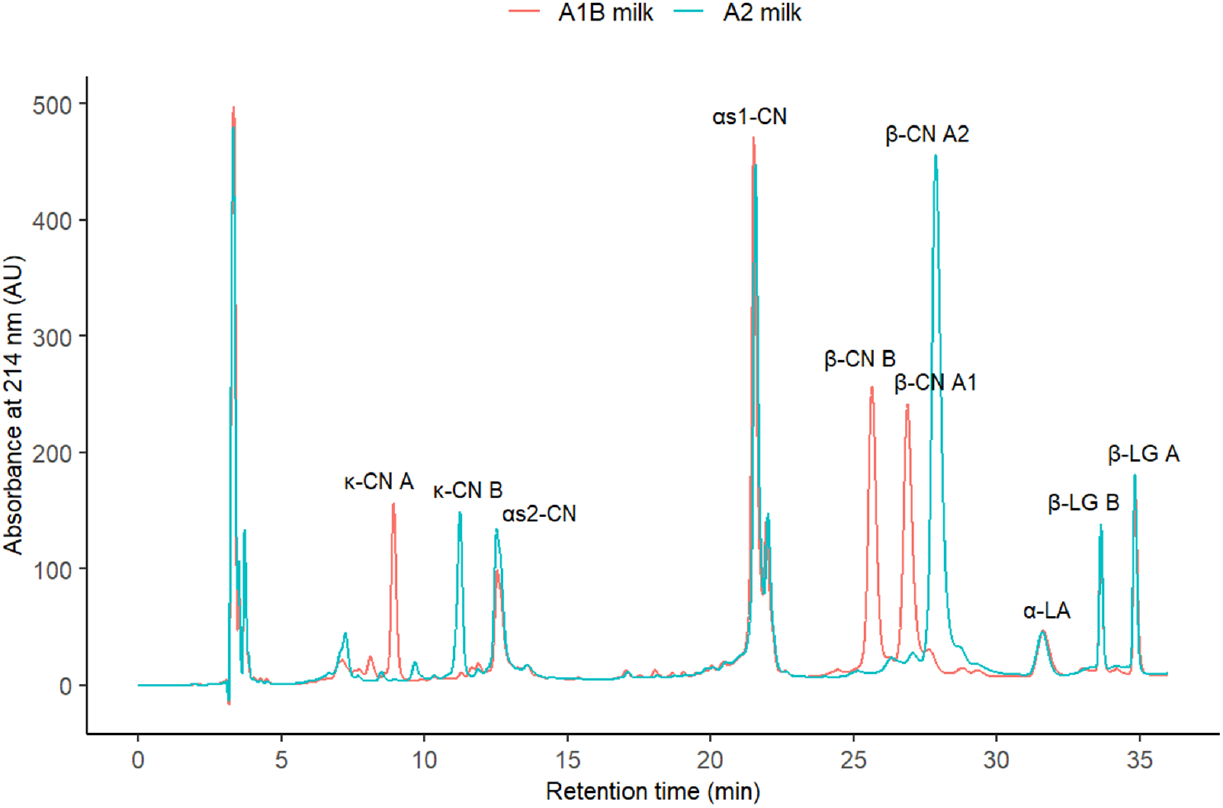
Representative RP-HPLC chromatograms of protein fractions from A1B and A2 milk. Under identical analytical conditions, RP-HPLC analysis reveals distinct elution profiles for β-CN A1, A2, and B variants. Major peaks are annotated: κ-casein A (κ-CN A), κ-casein B (κ-CN B), αs2-casein (αs2-CN), αs1-casein (αs1-CN), β-CN B (β-CN B), β-casein A1 (β-CN A1), β-casein A2 (β-CN A2), α-lactalbumin (α-LA), β-lactoglobulin B (β-LG B), and β-lactoglobulin A (β-LG A). Unlabeled peaks correspond to minor proteins, co-eluting components, or baseline features.

Other features of A1B and A2 milk are reported in Table 2. Milk samples exhibited comparable titratable acidity, casein index, and content of total solids, fat, total protein, lactose, and casein. The pH value and total mineral content were significantly higher in A1B milk than in A2 milk (p < 0.05). Additionally, besides the expected differences in the relative proportion of β-CN genetic variants to total β-CN (p < 0.05), the proportion of β-CN to total casein was significantly higher in A1B milk (0.43) compared to the A2 sample (0.40) (Chisquared = 3.86, p < 0.050). The β-CN fraction in A1B milk consisted of 64% A1 and 36% B variants, whereas A2 milk contained exclusively the A2 variant form. The higher proportion of β-CN observed in A1B milk was associated with a significantly lower αS1-CN-to-casein ratio, which was 0.36 in A1B milk compared to 0.39 in A2 milk (Chi-squared = 3.86, p < 0.050). These findings aligned with previous research, which reported significant effects of β-CN genotypes on the proportion of β-CN and αS1-CN in casein [20, 28].

**Table 2.**
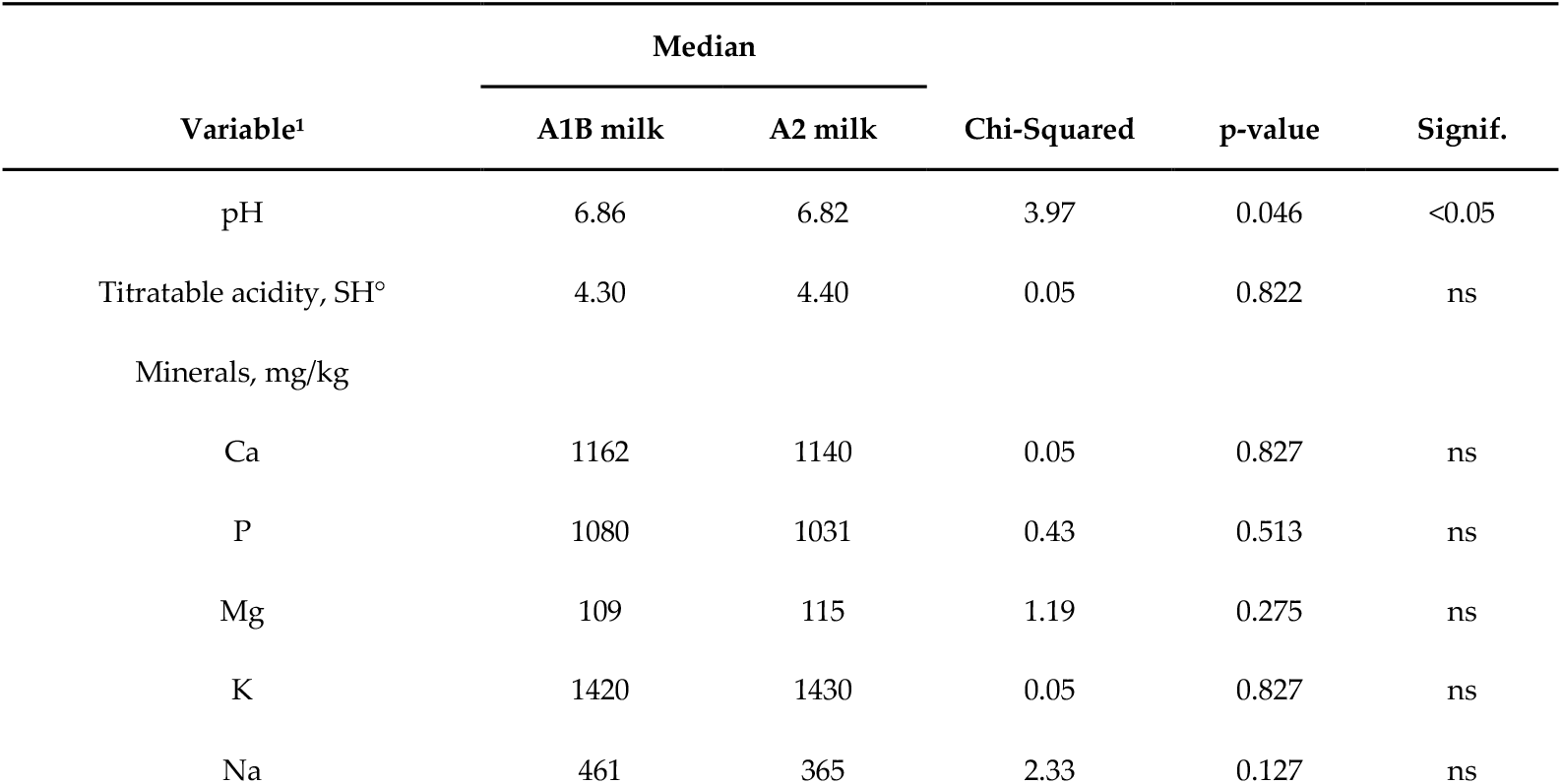
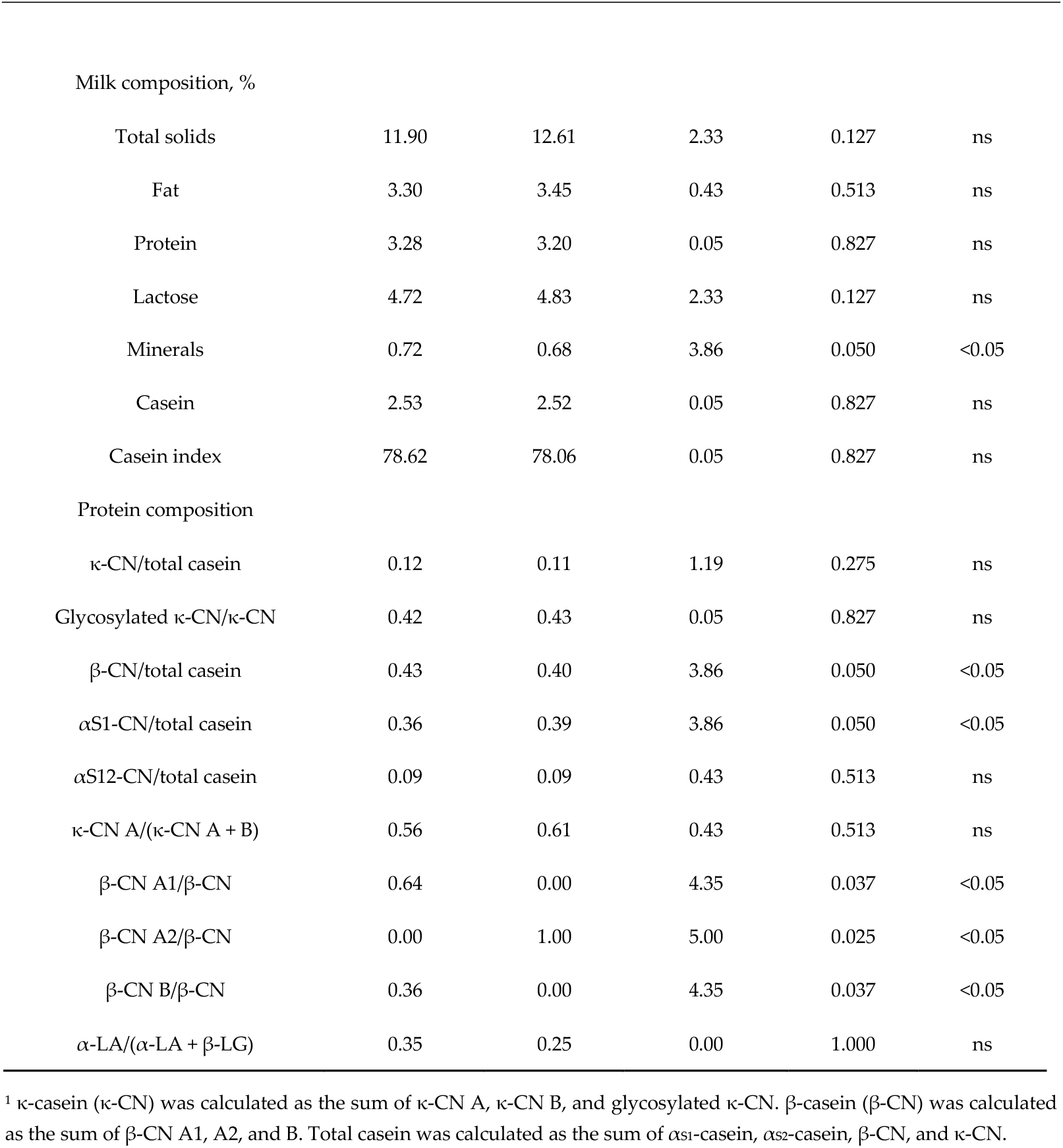
Median values (n = 3 per group) and results of Kruskal–Wallis tests for pH, titratable acidity, and chemical composition of A1B and A2 milk samples. Significant differences between milk types are indicated with their significance level; “ns” denotes non-significant differences.

| Variable<sup>1</sup> | Median |  | Chi-Squared | p-value | Signif. |
| --- | --- | --- | --- | --- | --- |
|  | A1B milk | A2 milk |  |  |  |
| pH | 6.86 | 6.82 | 3.97 | 0.046 | <0.05 |
| Titratable acidity, SH<sup>o</sup> | 4.30 | 4.40 | 0.05 | 0.822 | ns |
| Minerals, mg/kg |  |  |  |  |  |
| Ca | 1162 | 1140 | 0.05 | 0.827 | ns |
| P | 1080 | 1031 | 0.43 | 0.513 | ns |
| Mg | 109 | 115 | 1.19 | 0.275 | ns |
| K | 1420 | 1430 | 0.05 | 0.827 | ns |
| Na | 461 | 365 | 2.33 | 0.127 | ns |
| Milk composition, % |  |  |  |  |  |
| Total solids | 11.90 | 12.61 | 2.33 | 0.127 | ns |
| Fat | 3.30 | 3.45 | 0.43 | 0.513 | ns |
| Protein | 3.28 | 3.20 | 0.05 | 0.827 | ns |
| Lactose | 4.72 | 4.83 | 2.33 | 0.127 | ns |
| Minerals | 0.72 | 0.68 | 3.86 | 0.050 | <0.05 |
| Casein | 2.53 | 2.52 | 0.05 | 0.827 | ns |
| Casein index | 78.62 | 78.06 | 0.05 | 0.827 | ns |
| Protein composition |  |  |  |  |  |
| κ-CN/total casein | 0.12 | 0.11 | 1.19 | 0.275 | ns |
| Glycosylated κ-CN/κ-CN | 0.42 | 0.43 | 0.05 | 0.827 | ns |
| β-CN/total casein | 0.43 | 0.40 | 3.86 | 0.050 | <0.05 |
| αS1-CN/total casein | 0.36 | 0.39 | 3.86 | 0.050 | <0.05 |
| αS12-CN/total casein | 0.09 | 0.09 | 0.43 | 0.513 | ns |
| κ-CN A/(κ-CN A + B) | 0.56 | 0.61 | 0.43 | 0.513 | ns |
| β-CN A1/β-CN | 0.64 | 0.00 | 4.35 | 0.037 | <0.05 |
| β-CN A2/β-CN | 0.00 | 1.00 | 5.00 | 0.025 | <0.05 |
| β-CN B/β-CN | 0.36 | 0.00 | 4.35 | 0.037 | <0.05 |
| α-LA/(α-LA + β-LG) | 0.35 | 0.25 | 0.00 | 1.000 | ns |

<sup>1</sup> κ-casein ( κ-CN) was calculated as the sum of κ-CN A, κ-CN B, and glycosylated κ-CN. β-casein ( β-CN) was calculated as the sum of β-CN A1, A2, and B. Total casein was calculated as the sum of αS1-casein, $\alpha$ S2-casein, β-CN, and κ-CN.

### 3.2 Effects of DA1B and DA2 on function, protein composition, and inflammatory profile of the leaky IEB

To study the impact of DA1B and DA2 on the leaky IEB, we first verified that the digests did not affect the intestinal cell viability. After the 24 h treatment with DA1B and DA2, cellular viability ranged between 88.9 ± 1.5 and 90.37 ± 1.6% of live cells with respect to total cells, indicating that the treatments were not cytotoxic. In the same experiments, samples constituted by the buffer and enzymes used for *in vitro* digestion were also tested without showing any differences from NT cells (88.90 ± 1.64% live cells).

Successively, TEER and intestinal paracellular permeability were measured to evaluate the effect of DA1B and DA2 on the functions of the leaky IEB. As reported in Figure 2a, in the untreated leaky co-culture (NT), TEER was 44.74 ± 3.66 Ω cm^2^. The administration of DA1B or DA2 did not significantly modify TEER (Figure 2a) and paracellular permeability (Figure 2b). Then, we studied whether DA1B and/or DA2 were able to protect leaky IEB. To this aim, the leaky co-culture was treated with LPS and a mixture of TNFα and IFN-γ (MIX) in the presence or absence of milk digests. The application of an inflammatory stimulus impaired the IEB, TEER decreased to 62.78 ± 9.42% of NT, and the permeability increased to 161.59 ± 10.49 % of NT, values that were not modified by the presence of the digests (Figure 2a and b).

**Figure 2.**
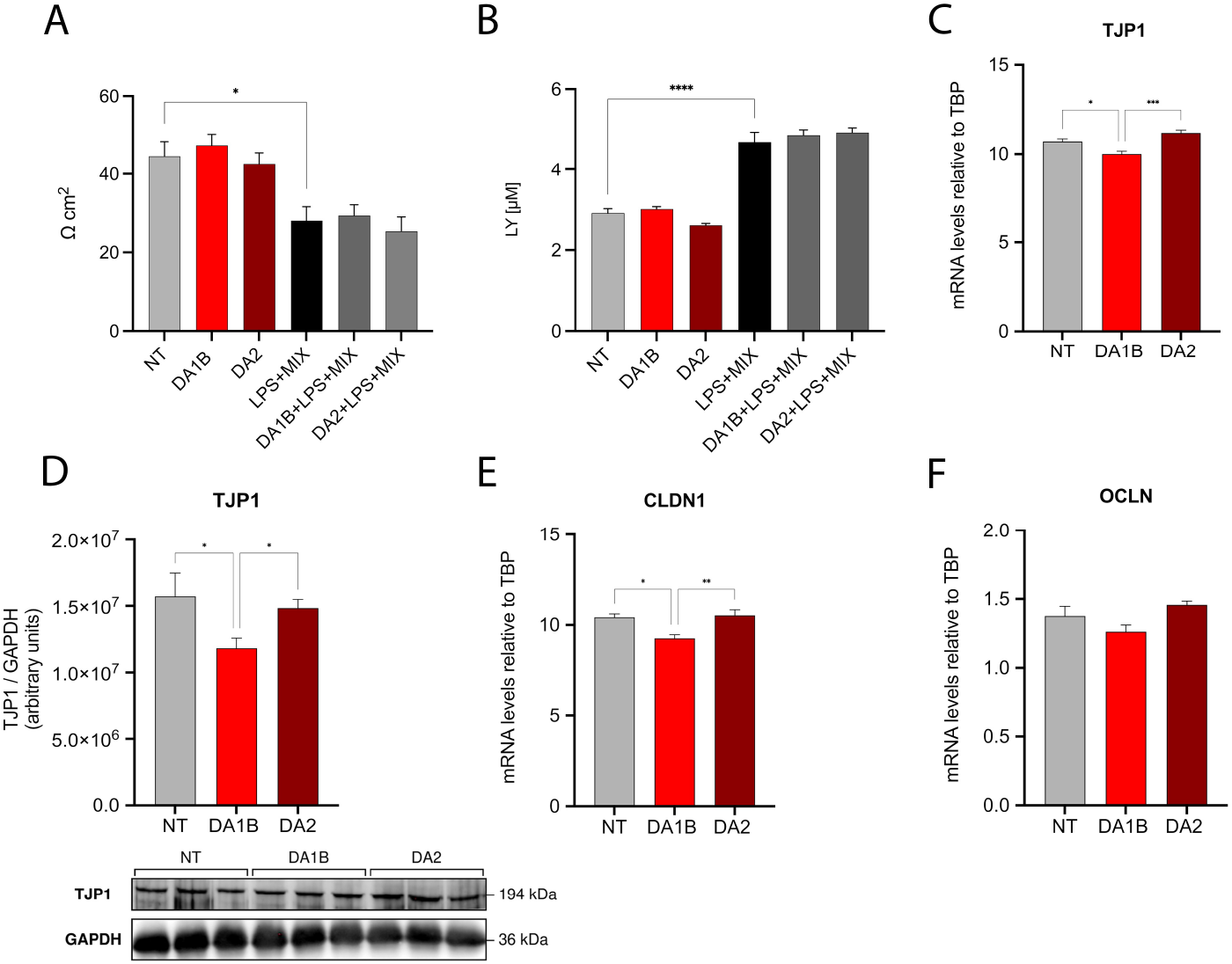
Effects of DA1B and DA2 on leaky IEB integrity. (**a**) TEER values expressed as Ω cm^2^ in leaky co-culture treated for 24 h with digests. (**b**) paracellular permeability to Lucifer Yellow expressed as [LY] in the basolateral compartment of the transwells in leaky Caco-2/HT-29 co-culture treated for 24 h with digests. (**c**), (**e)**, and (**f)**, relative gene expression levels of tight junction protein 1 (TJP1), claudin 1 (CLDN1), and occludin (OCLN) after 24 h exposure to DA1B and DA2. (**d)**, representative immunoblot and protein quantification of TJP1 in leaky Caco-2/HT-29 co-culture treated with DA1B and DA2 for 24 h. NT, untreated cells; DA1B, A1B milk digestate; DA2, A2 milk digestate; LPS, lipopolysaccharide; MIX, mixture of TNFα and IFN-γ; LY, Lucifer Yellow. (**a)** and (**b)** Two independent experiments with at least three technical replicates for each sample. In each sample, three TEER measurements were performed. Data show means ± SD. (**c**-**f)** Three independent experiments with three technical replicates for each experiment were performed; *p-value < 0.05; **p-value < 0.01; ****p-value < 0.0001. Data are presented as mean ± SD.

To deeply consider whether DA1B and DA2 can affect the protein composition of the leaky gut, we evaluated the expression of key genes involved in tight junctions, 24 h after exposure to the digests, compared to untreated cells. The gene expression of zonula occludens-1 (TJP1), as well as the protein level, showed a down-regulation when the leaky co-culture was treated with DA1B (Figure 2c and d). Claudin-1 (CLDN1) gene expression paralleled that of TJP1, while occludin (OCLN) was not affected (Figure 2e and f). DA2 did not affect the expression of any tight junction protein.

Since intestinal cells can release cytokines that regulate IEB homeostasis and modulate the inflammatory response, we studied whether the expression of interleukin-8 (IL8), TNFα, and IL6 was affected by DA1B and DA2. The IL8 gene expression was upregulated after 24 h of treatment with DA1B and DA2 (Figure 3a). The ELISA performed on cell culture supernatants confirmed that IL8 secretion was significantly increased after 24 h treatment, and no significant differences were observed between DA1B and DA2 treatments (Figure 3b). Notably, after 24 h treatment, DA1B digest induced a significant increase in TNFα gene expression compared with NT (Figure 3c). The IL6 expression remained unchanged (Figure 3d).

**Figure 3.**
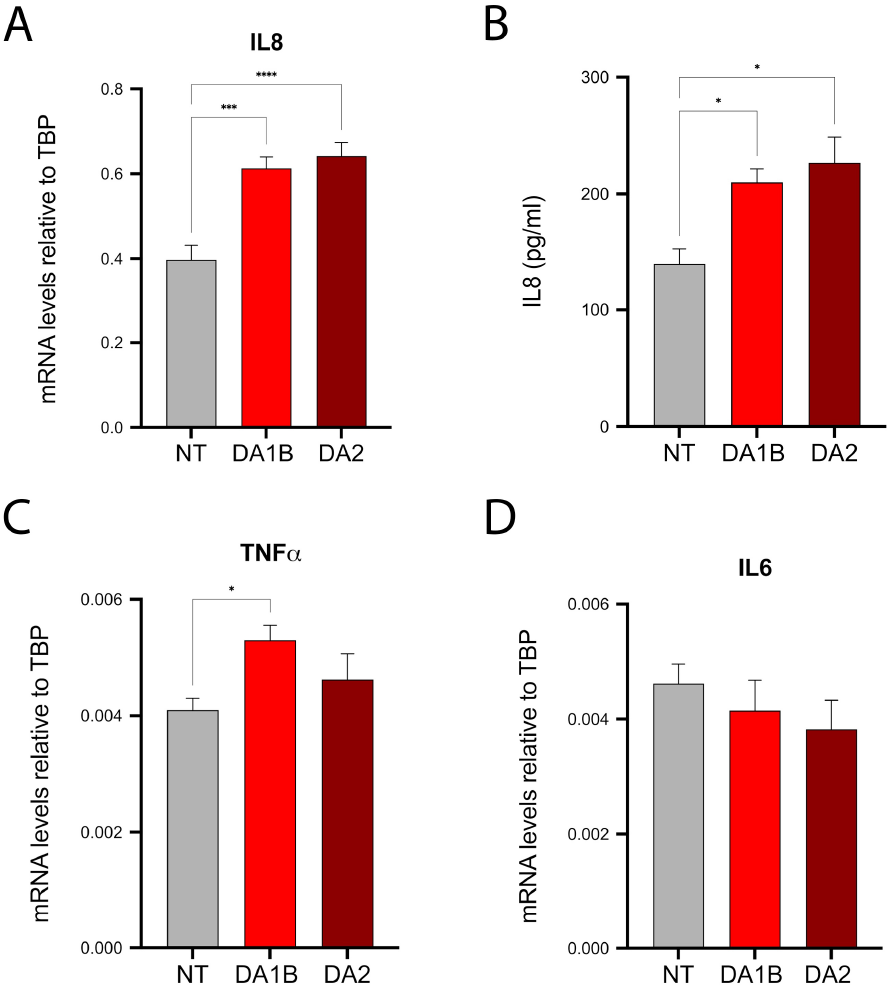
Effect of DA1B and DA2 on the inflammatory profile of leaky IEB. (**a**) and (**b**), IL8 relative gene expression levels and concentration measured by ELISA in the leaky co-culture supernatants after 24 h of DA1B and DA2 administration. (**c**) and (**d**), relative gene expression levels of TNFα and IL6 after 24 h exposure to DA1B and DA2 administered to leaky co-culture. NT, untreated cells; DA1B, A1B milk digest; DA2, A2 milk digest. Three independent experiments with three technical replicates for each experiment were performed; * p-value < 0.05; ***p-value < 0.001; ****p-value < 0.0001. Data are presented as mean ± SEM.

### 3.3. Effects of MA1B and MA2 on hMADS adipocytes

To assess the effects of A1B and A2 milk on human adipocytes, hMADS fully differentiated adipocytes (Figure 4a) were treated with MA1B and MA2 for 24 h, and the expression of inflammatory cytokines and adipokines was subsequently evaluated.

**Figure 4.**
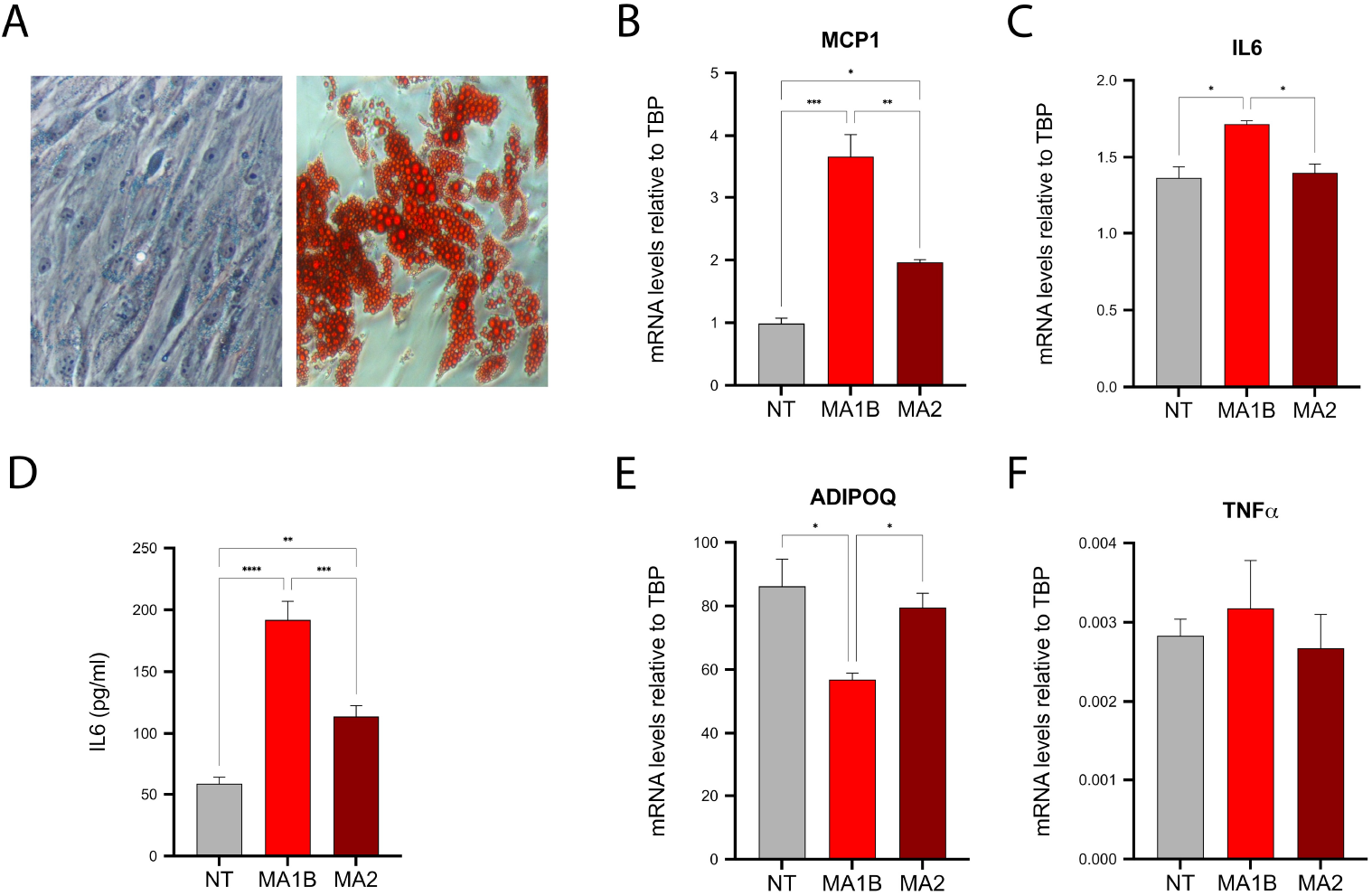
Effects of MA1B and MA2 on hMADS adipocytes. (**a**), light microscopy representative pictures of undifferentiated hMADS cells (day 0, left panel) and Oil Red O-stained differentiated hMADS adipocytes (day 13, right panel). Relative gene expression levels of MCP1 **(b**), IL6 (**c**), adiponectin (ADIPOQ) (**e**), and TNFα (**f**) in hMADS adipocytes after 24 h exposure to MA1B and MA2. Data are presented as mean ± SEM. (**d**), IL6 concentrations measured by ELISA in the culture supernatants of adipocytes after 24 h exposure to MA1B and MA2. NT, untreated cells; MA1B, digested and absorbed fractions derived from A1B digest; MA2 digested and absorbed fractions derived from A2 milk digest. Three independent experiments with three technical replicates for each experiment were performed; *p-value < 0.05; **p-value < 0.01; ***p-value < 0.001; ****p-value < 0.0001. Data are presented as mean ± SEM.

The results showed that MA1B significantly increased both the expression of MCP1, known to be involved in inflammatory cell infiltration in obese fat, and the proinflammatory cytokine IL6, compared to both NT and MA2-treated cells (Figure 4b and c). The IL6 mRNA levels were also confirmed by ELISA performed on cell culture supernatants. Both MA1B and MA2 induced IL6 release into the supernatant, although MA2 elicited a significantly lower response (Figure 4d). Conversely, MA1B significantly reduced the expression of the insulin-sensitizing hormone adiponectin (ADIPOQ, Figure 4e), while the expression levels of TNFα remained unchanged (Figure 4f). Collectively, these data suggest that MA2 may have a lower inflammatory potential than MA1B in adipose cells, with a less pronounced effect.

## 4. Discussion

To the best of our knowledge, this study represents the first *in vitro* study dealing with the effects of A2 milk on leaky gut syndrome in correlation with obesity.

Previous studies have explored the effects of the β-CN variants A1 and A2 on gut health. Although both are naturally occurring in bovine milk, several clinical and preclinical studies suggested that A1 β-CN may be associated with gastrointestinal discomfort, low-grade inflammation, and increased abdominal pain, whereas the consumption of A2 milk was hypothesized to be protective [29, 30].

Recently, the work of Guantario et al. [16] in an animal model of ageing mice demonstrated a positive role of A2 milk on gut immunology and morphology. Wang et al. [17] demonstrated that consumption of A2 milk was associated with a minor gastro intestinal discomfort in middle and old aged group of human adults. These two studies share the same intestinal model characterized by a higher permeability, i.e., leaky gut, which represents both a specific underlying condition for the onset of pathologies such as IBS, celiac diseases, diabetes, obesity, and gut microbiome dysbiosis [15], but also of the physiological senescence of intestinal cells [31].

Our study on an *in vitro* model of aged leaky gut [23] agrees with results reported in experiments in which 3-week-old Balb/c mice were fed with a diet supplemented with bovine milk containing A1 and A2 variants of β-CN or A2 type β-CN, without showing any direct variation of the intestinal permeability, either in the absence or the presence of inflammation [32]. Moreover, the same study reported the induction of a gut inflammatory response following the consumption of A1 β-CN and not of the A2-type variant.

In this regard, the work of Aitchison et al. [33] first reported that gene expression of inflammatory marker IL8 in Caco-2 cells with and without stimulation by IL1ß increased at the same level after administration of *in vitro* digested A1-like or A2-like β-CN, a result reproduced even in our leaky co-culture, by using DA1B and DA2. Nevertheless, DA1B increased TNFα, while DA2 did not, indicating a minor inflammatory state compared to treatments with digests of A1B milk.

To better elucidate these results, we also explored a possible effect on the leaky IEB integrity as it concerns the molecular composition of the tight junctions. The relation between nutrition, especially milk protein components, and IEB permeability has already been demonstrated in the case of CLDN1 expression and function [34], as well as of TJP1 level [35]. Under our experimental conditions, only DA1B administration decreases both CLDN1 and TJP1 expression level with respect to NT cells. Occludin, an IEB protein not modified in leaky aged co-culture [23], was unaffected by both treatments, once again indicative of a non-specific perturbation of permeability, already reported for casein and its peptides [36].

Conversely, there is a limited literature focusing on the systemic metabolic effects of milk, particularly in relation to body metabolism, obesity, and adipose tissue biology. *In vitro* studies have shown that milk can promote adipocyte differentiation in murine 3T3-L1 preadipocytes [37] and that milk-derived exosomes are particularly enriched in microRNAs associated with adipogenesis [38]. Notably, bovine milk exosomes have been shown to promote the expression of thermogenic and mitochondrial-related genes in both 3T3-L1 adipocytes and hADSCs [39]. Consistently, *in vivo* oral administration of milk exosomes in high-fat diet-fed mice resulted in reduced body weight. These effects were largely attributed to the bovine-specific miR-11987, which promotes the browning of white adipocytes, suggesting a potential role for milk-derived exosomes in the management of obesity and metabolic syndrome [39].

Given the well-established association between obesity, chronic inflammation, and insulin resistance, largely driven by macrophage infiltration in adipose tissue [40, 41], we investigated whether intestinal digested and absorbed fractions (MA1B and MA2) of milk samples exert differential effects on hMADS adipocytes, a widely accepted model of human white adipocyte [42].

Our data showed that the MA1B significantly increased the expression of MCP1, a key chemokine involved in the recruitment of inflammatory cells in obese fat [43], and IL6, a pro-inflammatory cytokine. Moreover, MA1B reduced the expression of ADPOQ, a well-known adipokine with insulin-sensitizing properties that suppresses NF-κB signaling and promotes metabolic homeostasis [44]. Conversely, MA2 maintained lower expression levels of MCP1 and IL6, while preserving adiponectin expression. Since elevated levels of MCP1 and IL6 are hallmark features of adipose tissue inflammation and contribute to obesity-induced insulin resistance [30, 45], the ability of MA2 digests to sustain ADPOQ levels further supports their protective role.

Collectively, these findings suggest that the *in vitro* digested and absorbed fraction of A2 milk may help to ameliorate the *in vitro* leaky IEB integrity, exhibit a lower inflammatory potential, and have a protective effect on obesity-related adipose tissue inflammation.

It is important to underline that although the experimentation reported used the *in vitro* cell models, we used milk rather than A1B and A2 purified proteins; thus, the whole food matrix and the interactions among the components can be responsible for the results described here, and for the differences obtained with other studies. In this context, a limitation of the present study could lie in other compositional differences observed in A1B and A2 milk samples. Based on this, the current experimental design did not allow us to determine with certainty whether the biological responses observed are attributable exclusively to β-CN polymorphisms or somewhat influenced by other factors. Nonetheless, the differences in pH and mineral content, lower β-CN-to-total casein ratio, and proportion of αS1-casein reflect inherent biological consequences of the underlying protein polymorphisms [20, 28]. Therefore, they represent structural features of A2 milk rather than a controllable source of variation. Notwithstanding, the intestinal *in vitro* model adopted here is a limitation, as it does not account for the presence of the gut microbiota, which plays an important role in the pathophysiology of leaky gut, obesity-related inflammation, and the metabolism of the two β-CN isoforms [46].

## 5. Conclusions

Significant differences between A1B and A2 milk samples did not emerge as it concerns *in vitro* leaky IEB integrity. The A2 milk exhibited a lower inflammatory potential in leaky gut, likely supporting the evidence that A2 milk can alleviate preexisting leaky gut discomfort.

This study is the first to establish a link between the effects of A2 milk, leaky gut syndrome, and obesity, suggesting a gut–adipose axis involvement. Future *in vitro* studies need to be designed taking into account all the possible influencing factors, for example BCM-7 peptide formation, microbiota changes, and/or their metabolites, all playing a role in the modulation of barrier function and permeability, a first step relating gut to adipose tissue, inflammation to obesity. In this context, dairy foods and their derivatives could play an important function to maintain the IEB integrity, ensuring positive effects on the target organs as well.

## Author Contributions

JP and AF: conception and design, data analysis and interpretation and manuscript writing. PB, ES, SC and CG: performance of experiments and figure preparation. IDN and VB: data analysis, interpretation and manuscript writing. AG and SC criticism during the study, revision of the manuscript and financial support. All authors approved the final version of the manuscript.

## Data Availability Statement

The raw data supporting the conclusions of this article will be made available by the authors on request.

## Acknowledgements

The authors are very grateful to Roberto Di Mulo, owner of the farm *Angolo di Paradiso* (Fermo, Italy), for providing milk samples and to Dr Christian Dani for providing hMADS cells.

**The Graphical abstract was partially created in** http://BioRender.com

## Abbreviations

The following abbreviations are used in this manuscript:

A1B: milk milk containing β-CN A1 and B variants
A2: milk milk containing β-CN A2 variant
β-CN: β-casein
BCM-7: β-casomorphin-7
DA1B: A1B in vitro milk digest
DA2: A2 in vitro milk digest
FA: fatty acid
hMADS: human multipotent adipose-derived stem cells
IEB: intestinal epithelial barrier
LY: lucifer yellow
MA1B: digested and absorbed fractions derived from A1B milk digest
MA2: digested and absorbed fractions derived from A2 milk digest
SCC: somatic cell count
SGID: static gastrointestinal digestion
TEER: transepithelial electrical resistance

